# Analyses of the Autism-associated Neuroligin-3 R451C Mutation in Human Neurons Reveals a Gain-of-Function Synaptic Mechanism

**DOI:** 10.1101/2021.12.07.471501

**Authors:** Le Wang, Vincent R. Mirabella, Rujia Dai, Xiao Su, Ranjie Xu, Azadeh Jadali, Matteo Bernabucci, Ishnoor Singh, Yu Chen, Jianghua Tian, Peng Jiang, Kevin Y. Kwan, ChangHui Pak, Chunyu Liu, Davide Comoletti, Ronald P. Hart, Chao Chen, Thomas C. Südhof, Zhiping P. Pang

**Author notes:** Equal contribution (Le Wang, Vincent R. Mirabella). Co-corresponding authors (Chao Chen), Thomas C. Südhof (Thomas C. Südhof) or Zhiping Pang (Zhiping Pang).

## Abstract

Mutations in many synaptic genes are associated with autism spectrum disorders (ASDs), suggesting that synaptic dysfunction is a key driver of ASD pathogenesis. Among these mutations, the R451C-substitution in the *NLGN3* gene that encodes the postsynaptic adhesion molecule Neuroligin-3 is noteworthy because it was the first specific mutation linked to ASDs. In mice, the corresponding *Nlgn3* R451C-knockin mutation recapitulates social interaction deficits of ASD patients and produces synaptic abnormalities, but the impact of the *NLGN3* R451C-mutation on human neurons has not been investigated. Here, we generated human knock-in neurons with the *NLGN3* R451C-mutation. Strikingly, analyses of *NLGN3* R451C-mutant neurons revealed that the R451C-mutation decreased *NLGN3* protein levels but enhanced the strength of excitatory synapses without affecting inhibitory synapses. No significant cell death and endoplasmic reticulum stress were detected. Importantly, the augmentation of excitatory transmission was confirmed *in vivo* with human neurons transplanted into mouse forebrain.

Using single-cell RNA-seq experiments with co-cultured excitatory and inhibitory *NLGN3* R451C-mutant neurons, we identified differentially expressed genes in relatively mature human neurons that corresponded to synaptic gene expression networks. Moreover, gene ontology and enrichment analyses revealed convergent gene networks associated with ASDs and other mental disorders. Our findings suggest that the *NLGN3* R451C-mutation induces a gain-of-function enhancement in excitatory synaptic transmission that may contribute to the pathophysiology of ASDs.

## Introduction

Autism spectrum disorders (ASDs) are characterized by social interaction deficits, repetitive patterns of behavior, and language impairments, and exhibit a 60%-90% genetic heritability^1-4^. Altered neuronal connectivity and synaptic dysfunction are hypothesized to underlie the pathophysiology of ASDs, similar to that of other neurodevelopmental disorders, including schizophrenia and intellectual disabilities^5^. Many genes encoding synaptic proteins were associated with ASDs and other neurodevelopmental disorders, including the genes encoding neuroligins (NLGNs) and neurexins (NRXNs) that are adhesion molecules which control the specification of synapse properties^6, 7^.

The first gene mutation that was linked to ASDs was a mutation of the highly conserved arginine-451 to cysteine in the *NLGN3* gene (*NLGN3* R451C-mutation)^8^. The functional impact of *NLGN3* R451C-mutation has been intensely studied in heterologous expression systems^9-11^ and in *Nlgn3* R451C-knockin mice^12, 13^. *Nlgn3* R451C-knockin mice showed that the R451C-mutation diminishes surface trafficking of NLGN3 protein, resulting in decreased levels of NLGN3^11, 13, 14^. Despite the reduction of *Nlgn3* expression in *Nlgn3* R451C-knockin mice, inhibitory synaptic transmission in the cortex^12^ and excitatory synaptic transmission in hippocampal pyramidal cells ^13^ and cerebellar Purkinje cells are enhanced^14^. *Nlgn3* R451C-knockin mice displayed impaired social interactions^12, 15^, a core symptom of ASDs, but surprisingly exhibited an increased spatial learning ability^12^ as well as enhanced repetitive behaviors^16^. Although another independently generated line of *Nlgn3* R451C-mutant mice revealed minimal aberrant behavioral phenotypes^17^, both strains of mutant mice exhibited region-specific synaptic deficits^13^. However, none of the synaptic changes and behavioral abnormalities observed in *Nlgn3* R451C-mice were detected in *Nlgn3* knockout (KO) mice^12, 13^. These findings suggested a gain-of-function mechanism of the *NLGN3* R451C-mutation that is likely cell-type dependent^12, 18^. Indeed, paired recordings in in the hippocampus revealed that the *Nlgn3* R451C-mutation causes a gain-of-function phenotype in one type of synapse that is different from the *Nlgn3* KO, and a loss-of-function phenotype in another synapses due to the decreased Nlgn3 proteins levels that is the same as that of the *Nlgn3* KO^19^. Although thus detailed information of the effect of the *Nlgn3* R451C-mutation exists for mice, no data are available for the effect of the *NLGN3* R451C-mutation in human neurons.

The development of efficient methods for generating neurons from human pluripotent stem cells ^20-23^ has provided an opportunity to use human neurons form modeling neuropsychiatric disorders^24-27^. Notably, mutations in the *NRXN*1^28, 29^ and in *NLGN4*^30, 31^ genes in human neurons have uncovered intriguing functional changes that appear to be specific to human neurons.

These studies provided important insights into how *NRXN1* and *NLGN4* mutations cause a synaptic dysfunction that could underlie the pathophysiology of ASDs. However, no such information is available for the *NLGN3* R451C-mutation. Therefore, we have now generated human embryonic stem (ES) cell lines with a *NLGN3* R451C-mutation, which enabled us to dissect the functional impact of the *NLGN3* R451C-mutation in human neurons. Analysis of R451C-mutant excitatory and inhibitory human induced neurons (iNs) revealed a reduction in NLGN3 expression, and an enhanced excitatory synaptic strength. No significant cell death and endoplasmic reticulum (ER) stress was found to be associated with the R451C-mutation. When R451C-mutant and control iNs were grafted into the mouse brain, a similar augmentation of excitatory synaptic transmission was observed in *NLGN3* R451C-mutant human iNs. Single-cell RNA-seq of co-cultured excitatory and inhibitory iNs revealed that the *NLGN3* R451C-mutation causes differential expression gene (DEG) changes in excitatory neuronal gene networks related to synaptic transmission. These DEGs were found to be associated with ASD, schizophrenia, and bipolar disorders but unrelated to major depressive disorder and obesity.

Our findings suggest that the *NLGN3* R451C-mutation dramatically impacts excitatory synaptic transmission in human neurons, thereby triggering changes in overall network properties that may be related to mental disorders.

## Methods details

### Cell Culture

Human H1 ES cells (WA01 WiCell Research Institute, Inc.) were cultured in 37 °C, 5% CO_2_ on Matrigel® Matrix (Corning Life Sciences)-coated plates in mTeSR1 medium (Stem Cell Technologies) and maintained as described previously^32^.

### Animals

All animal work was performed without gender bias under the Institutional Animal Care and Use Committee (IACUC) protocol approved by Rutgers University IACUC Committee.*Rag*^*2-/-*^ immunodeficient mice (JAX# 017708,) were used for cell transplantation. Wildtype C57BL/6J mice at postnatal day 0-4 (P0-4) were used for glial culture. Briefly, P0-P3 mouse cortex were digested with papain, 1 μM Ca^2+^, 0.5 μM EDTA solution for 15 min at 37 °C. Cells were passaged twice to eliminate mouse neuron cells and cultured with 10% fetal bovine serum in DMEM (Invitrogen).

### Gene Targeting

R451C-mutant ES cell lines were generated using CRISPR/Cas9 genome knock in^33^. Briefly, to target the *NLGN3* locus of the H1 ES cells, sgRNA designed from Optimized CRISPR Design Tool (http://crispr.mit.edu/) PX459 vector expressing Cas9 (Addgene #62988) and single stranded oligodeoxynucleotide (ssODN) were transfected using the Lipofectamine 3000 reagent (ThermoFisher Scientific, L300015). The ssODN contained 140 base pairs of homology arms flanking the mutation site carrying mutations for R451C, a Mul1 restriction enzyme site for screening, along with mutated PAM sequence. Individual clones were hand-picked for expansion and screening by PCR and sequencing. The two homozygous R451C knock-in clones were further subcloned before expansion and freezing.

### Lentiviral generation

Lentiviruses were produced as described^34, 35^ in HEK 293 FT cells (ATCC) by co-transfection with 3rd generation helper lentivirus plasmid (pMDLg/pRRE, VsVG and pRSV-REV) using calcium phosphate transfections. Lentiviral particles were collected in mTeSR1 medium, aliquoted, and stored in -80 °C. The following lentivirus constructs were used: (1) FUW-TetO-Ngn2-P2A-puromycin; (2) FUW-TetO-Ascl1-T2A-puromycin; (3) FUW-TetO-Dlx2-IRES-hygromycin; (4) FUW-rtTA, (5) L309-mCherry, and (6) FUW-Venus.

### Generation of iN cells

Excitatory- (i.e. Ngn2-iNs) and inhibitory-human iNs (AD-iNs) were generated as described ^36, 37^. Briefly, ESCs were dissociated by Accutase (Stem Cell Technologies) and plated in six-well plate coated with Matrigel® Matrix (Corning Life Sciences) and mTeSR1 containing RHO/ROCK pathway inhibitor Y-27632. At the same time of plating, lentivirus cocktails were prepared as described above (200 μl/well of six-well plate) was added. On day 1, the culture medium was replaced with Neurobasal medium (Gibco) containing doxycycline (2 μg/ml), and doxycycline was maintained in medium for ∼2 weeks. On days 2 and 3, infected cells were selected by puromycin and hygromycin (1 μg/ml). On day 4, mouse glia cells were plated on Matrigel-coated coverslips (5×10^4^ cells/single well of 24 well plate). On day 5, AD-iNs: Ngn2-iNs at 6:4 ratio (total of 2 ×10^5^ cells/well) were mixed and plated on a monolayer of mouse glial cells. On Day 5, media change occurred using Neurobasal medium containing growth factors (BDNF, GDNF, NT3, all at 10 ng/ml), B27 and 1% GlutaMax. Cytosine β-D-arabinofuranoside (AraC, 2 μM) was added in the medium to stop glial cell division. Cultures were used for experiments after 6 weeks.

### Cell transplantation

R451C-mutant and control Ngn2-iNs (14 days after induction) were dissociated and mixed at 1:1 ratio (1×10^5^ cells per μl in PBS). As described previously ^38, 39^, the cells were then stereotaxically (ML: ±1.0 mm, AP: -2.0 mm, and DV: -1.2 and -1.5 mm) injected into the brains of P0 to P3 Rag^2 -/-^ immunodeficient mice. The pups were weaned at P21 and were kept up to 6 weeks before characterization. We did not observe any tumor formation of the transplanted cells in any of the animals.

### Immunofluorescence experiments and analysis

Primary antibodies were diluted in blocking buffer (1% goat serum, 0.25% Triton X-100 and 1% BSA) at 4 °C and incubated overnight. Cultures were washed 3 times with blocking buffer and then incubated with secondary antibody (1: 500) at RT for 1h. Following 3x DPBS washes, coverslips were mounted onto glass microscope slides with mounting medium containing DAPI. The following antibodies were used: Oct4 (Millipore Sigma MAB4401, 1:2,000), Tra-1-60 (Millipore Sigma MAB4360, 1:1,000), Map2 (Abcam, AB5392, 1:1,000), Synapsin (Rabbit, 1:3,000, E028), VGAT (Millipore Sigma AB5062P, 1:500), Tuj1 (Sysy, 1:1000), Calnexin (Enzo ADI-SPA-860-D, 1:200), Calreticulin (Enzo ADI-SPA-600-D, 1:200) and Caspase-3 (Cell Signaling #9661, 1:200). At least three independent culture batches were used for all experiments.

#### Synaptic puncta density analysis

immunofluorescence was performed on iNs at day 42 with dendrite marker MAP2 and synaptic markers Synapsin-1 and VGAT. Confocal images were taken using a Zeiss LSM700 laser-scanning confocal microscope with a 63x objective. All images were acquired using Z stack maximal projection in Zeiss Zen blue software. Correlated synaptic puncta size and intensity were quantified by *Intellicount* as reported previously ^40^. Puncta density and primary processes were blindly counted and quantified.

#### ER structure analysis

Immunofluorescence was performed on iNs at day 42 with dendrite marker MAP2, and ER markers Calnexin and Calreticulin. Images of iNs were obtained as z-projections on a Zeiss LSM700 laser-scanning confocal microscope using a 63x objective. Confocal images were analyzed by ImageJ ^41^, as reported previously^42^. Using the line tool of ImageJ, a 10 µm line was drawn from the nuclear envelope towards the periphery of each cell analyzed. Intensity and coefficient variation (C.V.) was calculated all along the 10 µm line. iN soma ER area and intensity was calculated by using the Image J ROI tool and normalized with MAP2.

### Western Blot analysis

For induced neuron culture, at day 40-50, the cultures were washed once with PBS. The proteins were directly collected by 2X SDS loading buffer (4% SDS, 125mM Tris Hcl (pH=6.8), 20% glycerol, 5% 2-mercaptoethanol and 0.01% bromophenol blue)^43^. Protein samples then were denatured by heating to 95 °C for 5 min. The denatured protein samples were allowed to cool down and equal amounts of samples (30 μl) were analyzed by 4-15% TGX gels (Biorad # 4561084DC) for sequential Western blots on the same membranes. The protein from the gel was then transferred to a 0.45 μm nitrocellulose membrane (Biorad #1620115) using the BIO-RAD transferring system. Nitrocellulose membrane was blocked in 5% nonfat milk for 1 hour at room temperature. After the blocking step, membranes were incubated with primary antibody overnight at 4 °C. The following day, the blots were washed three times with Tris-buffered saline with 0.1% Tween 20 (TBST) before the addition of horseradish peroxidase-conjugated secondary antibody at room temperature for 1 hour. All ER stress–related proteins referred to here were purchased from Cell Signaling Technology as an ER Stress Antibody Sampler Kit (CST, #9956, all 1:1000). Rabbit Nlgn3 (RRID: AB_2571813, Frontiers Institute, Japan, 1:1000)^44^, Rabbit Syntaxin (I378, Südhof laboratory, 1:5000). Protein bands were visualized by the addition of Clarity Western ECL Substrate (Biorad # 1705060). The density of the bands was quantified using ImageJ software. Beta-actin was measured as a loading control.

### Gene Expression Analyses

Total neuronal RNA from three independently generated batches of cultures were prepared using TRIzol ® Reagent (Thermo Fisher Scientific). Human-specific primers were used for *NLGN3, MAP2, Tuj1, VGAT, Vglut1, Vglut2, GABBR1, ACTB*, and PCR reaction conditions followed the manufacturer’s recommendations. Student’s t-test was used to compare grouped control and R451C means. Evaluation of qPCR was performed blind to the genotype. See **Supplemental Table 1** for a complete list of primers used in this study.

### Electrophysiology

#### Cell culture electrophysiology

Whole-cell patch-clamp electrophysiology for co-cultured iN cells was performed as described ^32^. For spontaneous postsynaptic current and current-clamp recordings, K-gluconate internal solution was used, which consisted of (in mM): 126 K-gluconate, 4 KCl, 10 HEPES, 0.05 EGTA, 4 ATP-magnesium, 0.3 GTP-sodium, and 10 phosphocreatine. Spontaneous postsynaptic currents were recorded at a holding potential of -70 mV in the presence of 20 μM CNQX or 50 μM picrotoxin (PTX) added in HEPES solution. For miniature postsynaptic current recording, Cs-based solution was used, which consisted of (in mM): 40 CsCl, 3.5 KCl, 10 HEPES, 0.05 EGTA, 90 K-gluconate, 1.8 NaCl, 1.7 MgCl_2_, 2 ATP-magnesium, 0.4 GTP-sodium, and 10 phosphocreatine. The recording external solution contained (in mM): 140 NaCl, 5 KCl, 10 HEPES, 2 CaCl_2_, 2 MgCl_2_, 10 Glucose, pH 7.4. Miniature postsynaptic currents were recorded at a holding potential of -70 mV in the presence of 20 μM CNQX or 50 μM PTX and 1 μM tetrodotoxin (TTX). All cell culture recordings were conducted at room temperature.

#### Brain slice recording

Mice were anesthetized and brains were quickly removed into ice-cold oxygenated artificial cerebrospinal fluid (ACSF) cutting solution (in mM): 50 sucrose, 2.5 KCl, 0.625 CaCl_2_, 1.2 MgCl_2_, 1.25 NaH_2_PO_4_, 25 NaHCO_3_, and 2.5 glucose). Coronal section slices at 300 µm were cut using a vibratome (VT 1200S; Leica). After 1 h recovery (33 °C) in ACSF (in mM)125 NaCl, 2.5 KCl, 2.5 CaCl_2_, 1.2 MgCl_2_, 1.25 NaH_2_PO_4_, 25 NaHCO_3_, and 2.5 glucose, slices were transferred to a recording chamber and perfused with ACSF at 30°C. Whole-cell patch-clamp recordings were performed as described ^45^.

Cells were excluded from analysis if series resistance (Rs) changed by more than 20% during the recording. In addition, any recordings where the access resistance was greater than 25 MΩ were also excluded from the analysis. Electrophysiological data were analyzed using Clampfit 10.5 (Molecular Devices).

### Visualization of the targeting gene expression

To visualize the spatiotemporal expression of the *NLGN1, NLGN2, NLGN3, NLGN4X*, and *NLGN4Y* in the human brain, we used the median value of these genes as the signature value to visualize in the GTEx v8 data and BrainSpan data. This analysis was performed using cerebroViz R package [https://github.com/ethanbahl/cerebroViz]^46^. We also tested the expression of *NLGN3* in the single cell data of ASD and visualized with heatmap using ggplot2 R package. All analysis was performed in R v3.6.

### Single-cell RNA-seq library preparation

For each cell line, two wells of a six-well plate were plated with mixed cultures at 2×10^6^ iNs and 2.5 × 10^5^ glia/well. On day 40, cells were dissociated with TrypLE (Thermo Fisher) for 5 min at 37°C, which preferentially releases neurons from glia. Cells were washed twice with 1X HBSS (Thermo Fisher), resuspended at 1,200 cells/μl in 1% Bovine Serum Albumin (BSA)/1X PBS and placed on ice. Single-cell cDNA libraries were generated using the Chromium Next GEM Single Cell 3’ GEM Reagent Kits v3.1 (10X Genomics). Approximately 2×10^4^ dissociated cells were loaded onto a cassette in the Chromium Controller with accompanying reagents to generate Gel Beads in Emulsions (GEMs). These were reverse transcribed to single-cell cDNA libraries before the oil emulsion was disrupted and cDNA was purified using Dynabeads MyOne Silane (Thermo Fisher). cDNA was amplified by PCR for 11 cycles and purified using SPRIselect reagent (Beckman-Coulter). cDNA fragmentation, A-tailing, and end repair was performed followed by adapter ligation for paired-end sequencing. An additional 12 cycles of PCR were performed to incorporate the sample index sequences and to amplify the libraries before purification and sequencing. Single cell RNA-seq libraries were sequenced by Psomagen® using the Illumina Novaseq platform.

### Single-cell gene expression data preprocessing

Raw reads were aligned with pooled mouse (mm10) and human (GRCh38) reference genomes and pre-processed using Cellranger software (v. 4.0.0, 10X Genomics). Analysis of reads mapped per cell identified >90% human cells, < 10% cell mouse cells (Fig. 3 A and S5). After loading cell x gene data into R using the Seurat (v. 4.0.2) package ^47, 48^, only cells expressing human genes were isolated for further analysis. After filtering (number of features between 200 and 10,000 per cell, less than 10% mitochondrial RNA), normalization, scaling, and PCA projection, the UMAP (Uniform Manifold Approximation and Projection) coordinates were calculated, and graph-based clustering was performed.

### Identifying differentially expressed genes (DEGs)

Differential gene expression analysis was performed by applying a Mann–Whitney U-test on three relatively most mature cell types (excitatory, NPY, and SST inhibitory neuron), comparing combined R451C samples with control. P-values were adjusted by Benjamini-Hochberg to a False Discovery Rate (q value). All graphs and analyses were generated and performed in R.

### SynGo annotation

For analysis of DEGs associated with synaptic function, GO analysis was performed using the SynGO^49^ website: https://www.syngoportal.org/.

### Cell-cell communication analysis

Cell communication analysis was conducted in the CellChat package (version 0.0.1)^50^. The library-size normalized scRNA-seq data were loaded into CellChat. The human ligand-receptor pairs in the database CellChatDB were used for calculating the cell interactions. The main steps for constructing cell-cell communication networks are as follows: 1) identifying over-expressed ligand or receptors in one cell type and their interactions based on scRNA-seq data, 2) calculating the probability of cell interactions at ligand-receptor pair level, 3) inferring the cell communication probability at the signal pathway level. Networks were constructed in the control cell line and R451C cell line separately. The comparison of the total number of interactions and strength (probability of interaction) between two cell lines was conducted with the function *compareInteractions*. Mann-Whitney U test was used to compare the strength of interactions mediated by pathways *NRXN* and *NEGR*.

### Protein-protein interaction (PPI) network construction

The Search Tool for the Retrieval of Interacting Genes/Proteins database (STRING v11.0)^51^ was used to construct PPI network. Given gene NLGN3 (protein NLGN3) and the lists of differentially expressed genes in NPY neurons, SST neurons, and excitatory neurons (adjusted p value<0.05), STRING can search the proteins that have interactions with NLGN3 and construct subnetworks. K-means was used for network clustering. The genes connected to NLGN3 were defined as NLGN3-PPI-genes.

### Overrepresentation analysis of psychiatric disorder-related genes

To explore whether DEGs may be part of the same network or pathway as known susceptibility genes of psychiatric disorders, we tested for enrichment between DEGs and disorder-associated genes. Fisher’s exact test was used in the enrichment test. Significance was assessed using Fisher’s exact test, followed by FDR-correction of p values and OR > 1. We collected the genes from multiple resources which were classified into 46 categories. The gene identifiers were converted to Ensembl Gene IDs in Gencode (version v19 hg19).

### Quantification and Statistical Analysis

All data are presented as the mean ± standard error of the mean (SEM), unless otherwise noted. All experiments were repeated on at least three biological replicates, with each replicate consisting of an independent infection/differentiation. Statistical significance (**, p<0.01; ***, p<0.001) was evaluated with the Kolmogorov-Smirnov-test (for cumulative probability plots) and Student’s t-test.

### Data availability

Single cell-seq data will be deposited and available on the NCBI Gene Expression Omnibus under accession number GSE180751.

## RESULTS

### *NLGN3* R451C-mutation affects excitatory synapse formation and increases the strength of excitatory synaptic transmission

We first examined the expression of *NLGN3* in the human brain by evaluating the region-wide expression of *NLGN* genes (*NLGN1-4*) in brain using the Genotype-Tissue Expression (GTEx) database^52^. *NLGN2* and *NLGN3* are both abundantly expressed in the human frontal lobe (FL) and cerebellum (CB) (**Supplemental Fig. 1a & b**). Using the BrainSpan human brain development database^53^, we found that *NLGN3* is present at high levels during late prenatal and early postnatal development (**Supplemental Fig. 1c & d**). Consistent with current ASD *de novo* variant genetic findings, ASD risk genes related to synaptic communication are highly expressed in the brain at perinatal stages and have unique expression patterns compared with other groups of risk genes^54^. *NLGN* genes are expressed in both glia and neurons^55, 56^. To verify the cell type-specific expression of *NLGN3* in the human brain, we examined the ASD patient single-cell dataset^57^ and found that the *NLGN3* gene is expressed in both neuronal (GABA and glutamate neurons) and glial cells (**Supplemental Fig. 1e**). Although the gene expression data thus suggest that *NLGN3* may have a function in both neurons and in glia, we decided to focus here on neurons because the goal of the current studies was to determine whether the well-documented role of NLGNs in murine synapses^6, 58, 59^ also applies to human neurons.

To study the potential role of the *NLGN3* R451C-mutation in human neurons, we used CRISPR-Cas9 to introduce the R451C-mutation into human H1 ES cells (**Fig. 1a&b**). We selected two subclones (clone A&B) and used the parental ES cell line at a similar passage number as wild-type (WT) control for morphological, functional, and genomic analyses. Note that since the *NLGN3* gene is X-chromosomal and H1 cells are male, only a single allele needed to be mutated. We confirmed the pluripotency of NGLN3 R451C-mutant ES cells by staining the cells for the stem-cell markers Tra-1-60 and OCT4 (**Fig. 1c**), and additionally excluded off-targets mutations by sequencing the genomic DNA at the top 10 predicted CRISPR off-target sites.

**Fig. 1.**
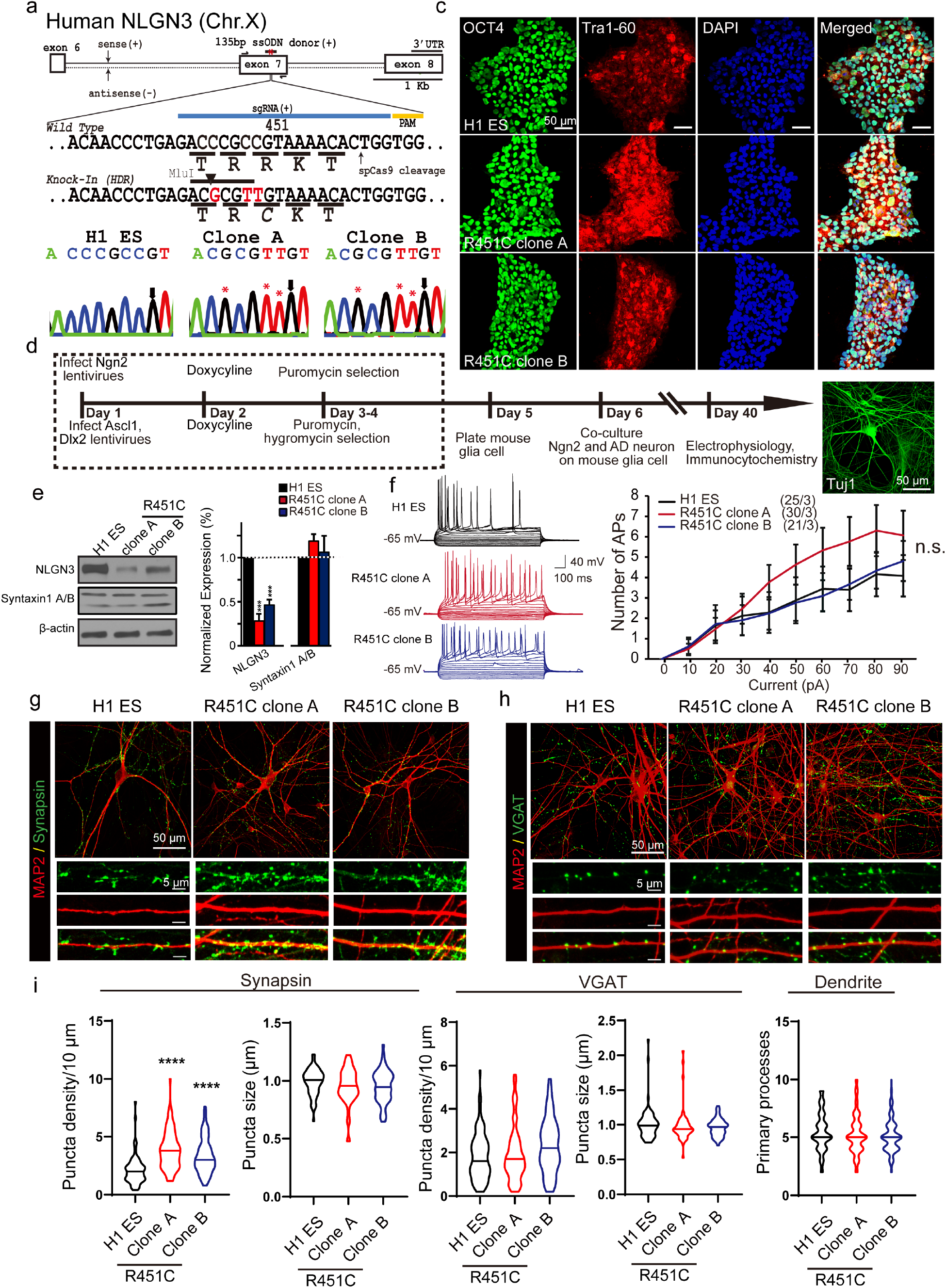
NLGN3 R451C mixed culture of induced neuronal (iN) cells. **a** NLGN3 R451C targeting strategy. The blue line indicates the sgRNA sequences, and the yellow line is the PAM. 135bp single-stranded oligodeoxynucleotide (ssODNs) is the base for direct homologous recombination of NLGN3 exon 7 genomic sequence. We inserted a Mlu1 restriction site for genotyping. **b** Confirmation of NLGN3 R451C knock-in cell lines by Sanger sequencing. * indicates mutated nucleotide. **c** Immunostaining of stem cell marker genes in control and converted ES cell lines. DAPI was used to visualize the cell nucleus. OCT4 (green) is expressed in the nucleus colocalized with DAPI, and Tra-160 (red) is expressed in the cell body. **d** Experimental design of *Ngn2* (GFP) and *Ascl1, Dlx2* (mCherry) iNs mixed culture paradigm. **e** Quantification of NLGN3 and Syntaxin protein expression levels in mixed culture of iN cells, N=3. **f** Representative traces and quantification plot of step current injection action potential firing number. **g** Representative images of control and R451C converted iNs total synapses immunolabeled with Synapsin (green) and MAP2 (red). **h** Representative images of control and R451C converted iNs inhibitory synapses immunolabeled with VGAT (green) and MAP2 (red). **i** Quantification of Synapsin/VGAT puncta density per 10 µm (n=90), puncta size (Synapsin, n=60; VGAT, n=70) and dendrite formation (n=130), statistical significance was assessed by Student’s t-test (^***^P < 0.001).

Using our well-characterized iN cell technology^35^, we converted human control and *NLGN3* R451C-mutant ES cells into excitatory (with Ngn2) and inhibitory (with Ascl1 and Dlx2) neurons^35-37^(**Fig. 1d**). We co-cultured the two types of neurons and performed protein and gene expression analysis. Consistent with what was found in the mouse model of the *Nlgn3* R451C mutation^12^, we observed that the R451C substitution caused a significant decrease in *Nlgn3* protein levels (∼40% of control levels), but not in the levels of Syntaxin-1 (**Fig 1e**). The *NLGN3* mRNA levels, however, were not changed by the R451C-knockin mutation as determined by quantitative RT-PCR (**Supplemental Fig. 2**). These data suggest that the *NLGN3* R451C-mutation destabilizes the *NLGN3* protein, consistent with previous studies on the mutant protein^9-13^.

To investigate whether *NLGN3* R451C-mutant neurons exhibit functional impairments, we next analyzed them by whole-cell patch-clamp electrophysiology and morphological measurements. We found that *NLGN3* R451C-mutant neurons have intrinsic electrical properties and fire action potentials in response to current injections similar to control neurons (**Fig. 1f, 2c-e**). However, neurons derived from both clones of the *NLGN3* R451C-mutant ES cells exhibited a significantly increased density of presynaptic Synapsin-positive puncta, suggesting an increased synapse number. In contrast, the density of inhibitory vGAT-positive synaptic puncta was not changed, indicating that only excitatory synapse numbers are increased (**Fig. 1g-i**). Moreover, the number of primary dendritic processes showed no significant changes (**Fig. 1i**), suggesting that the R451C-mutation does not affect dendritogenesis.

**Fig. 2.**
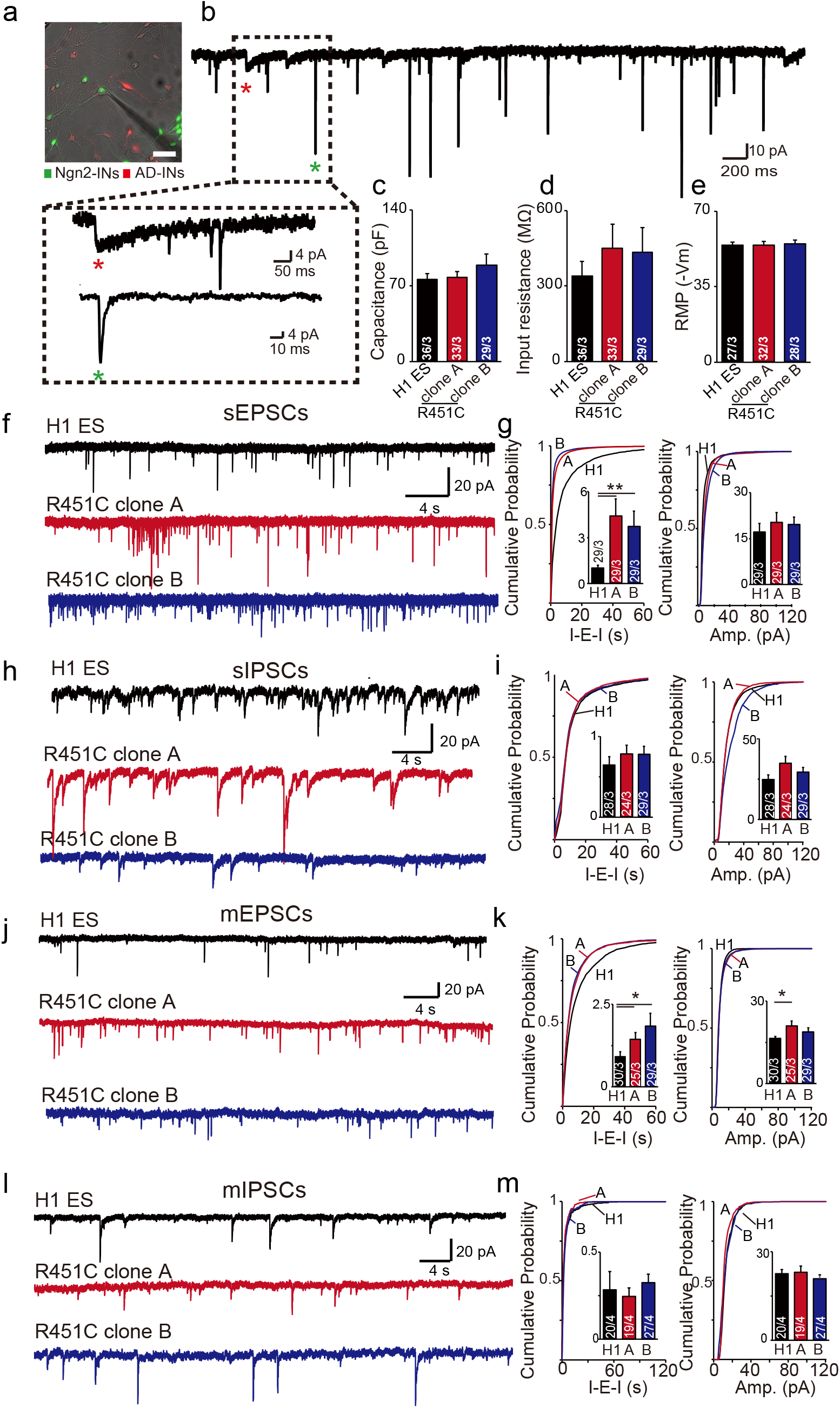
NLGN3 R451C increases excitatory neurotransmission in vitro. **a** Representative image of recording mixed culture iNs. Green cells are *Ngn2* iNs, red cells are *Ascl1* and *Dlx2* iNs. **b** Representative sPSC trace of mixed culture iNs. Green * indicate EPSC-like event, red ^*^ indicates IPSC-like event. **c-e** Intrinsic electrophysiology characterization of control and converted R451C iNs. Summary of passive membrane properties (capacitance, input resistance, resting membrane potential). **f** Representative traces of sEPSC. **g** Quantification of sEPSC frequency and amplitude. **h** Representative traces of sIPSCs. **i** Quantification of sIPSC frequency and amplitude. **j** Representative traces of mEPSC. **k** Quantification of mEPSC frequency and amplitude. **l** Representative traces of mIPSCs. **m** Quantification of mIPSC frequency and amplitude. Data are mean ± SEM; Number of cultured cells/ cultured batches are shown in bars. Statistical significance (^*^, p<0.05, ^**^, p<0.01) was evaluated with the Kolmogorov-Smirnov test (cumulative probability plots) and Student’s t-test (bar graphs).

Next, we investigated synaptic transmission using the co-cultured excitatory and inhibitory neurons (**Fig. 2a**). We simultaneously recorded spontaneous excitatory (sEPSCs) and inhibitory postsynaptic currents (sIPSCs) from *NLGN3* R451C-mutant and control neurons and distinguished between sEPSCs and sIPSCs based on their distinct kinetics^60^ (**Fig. 2b**). The *NLGN3* R451C-mutant neurons exhibited a large (∼3 fold) increase in the frequency, but not the amplitude, of sEPSCs without significant changes in sIPSCs (**Fig. 2f-i**). In the presence of tetrodotoxin (TTX), we detected a similar increase in miniature excitatory postsynaptic currents (mEPSCs) but not in miniature inhibitory postsynaptic currents (mIPSCs) in *NLGN3* R451C-mutant neurons compared to controls (**Fig. 2j-m**). Together, these results suggest that in human neurons, the *NLGN3* R451C-mutation enhances the strength of excitatory but not of inhibitory transmission at least in part by increasing synapse numbers.

### The *NLGN3* R451C-mutation is unlikely to cause ER stress

The R451C-mutation decreases the levels of *NLGN3* protein without lowering expression of the NLGN3 gene in human neurons (Fig. 1e, S2), similar to observations made in *Nlgn3* R451C-mutant knockin mice^12^. These observations prompted the hypothesis that the *NLGN3* R451C-mutation might impair neuronal function by inducing an ER stress response^14, 61^. Although it seems unlikely that ER stress would induce an increase in the number and strength of excitatory synapses, at least in cerebellar Purkinje cells it was suggested that an unfolded protein response (UPR) could enhance excitatory synaptic transmission^14^. These findings prompted us to investigate whether the *NLGN3* R451C-mutation induced ER stress or cell death in human neurons.

We first examined the morphology of the ER in human neurons. Staining of control and *NLGN3* R451C-mutant neurons showed that the signal of the ER marker calnexin did not show significant changes, whereas the calreticulin signal exhibited an increase in neurons derived from one but not from the other of the two *NLGN3* R451C-mutant ES cell clones that we analyzed (**Fig. 3a-l**). Moreover, quantitative immunoblotting for major ER stress markers, including Calnexin, Ero1-Lα, IRE1α, PDI and PERK, did not uncover consistent changes in the expression of these markers (**Fig. 3m & n**). Finally, we probed the neurons for evidence of cell death, but detected no significant change in the total number of neurons or the number of neurons that were stained by the apoptosis marker caspase-3, although there was a trend of a decrease in the former and of an increase in the latter (**Fig. 3o & p**). Together, these results suggest that the *NLGN3* R451C-mutation does not cause a major impairment of neurons by ER stress, although a low level of ER stress activation cannot be ruled out.

**Fig. 3.**
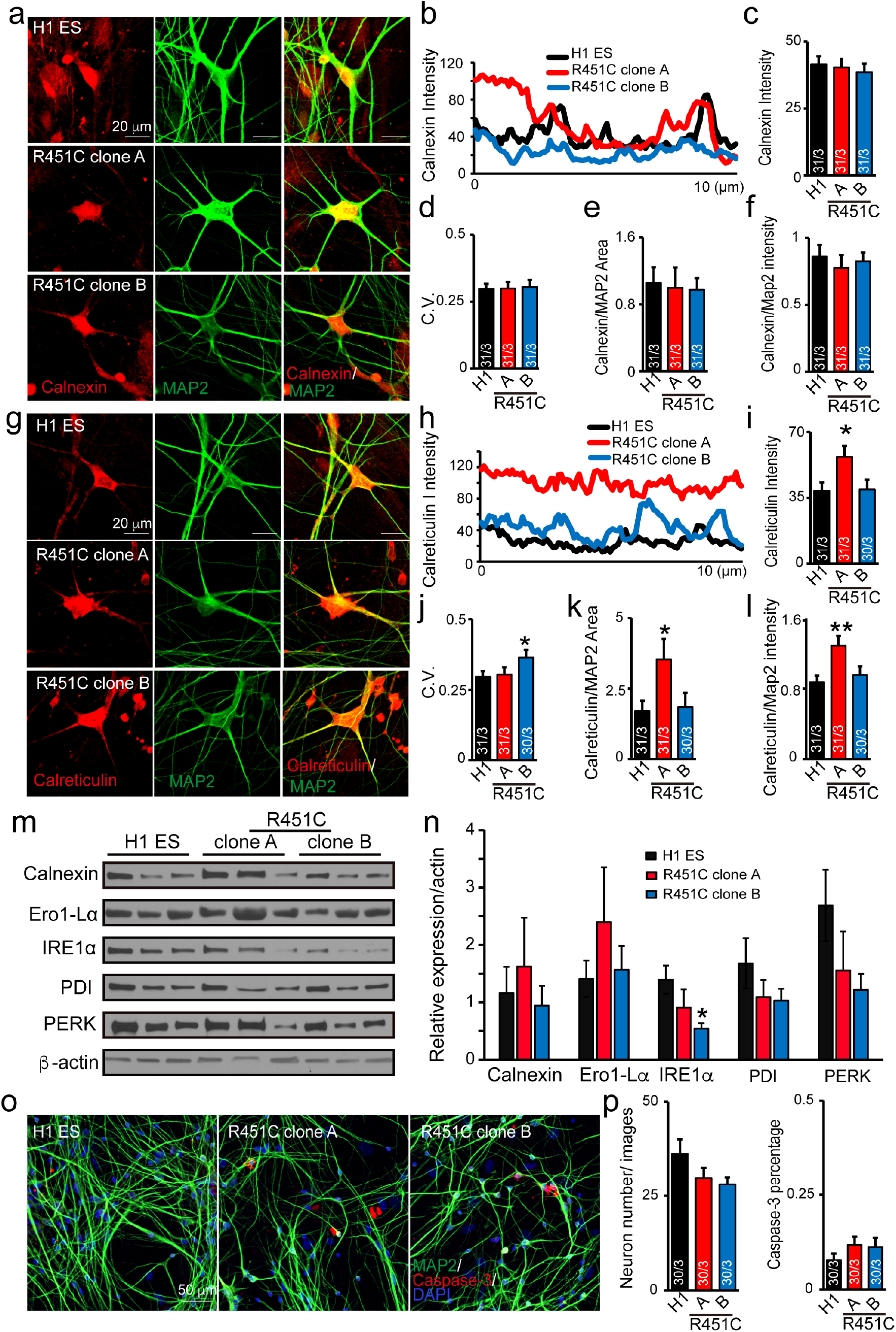
No major ER stress found in human neurons carrying NLGN3 R451C-mutation. **a** Representative image of Calnexin (red) and MAP2 (red) in R451C and control iNs. **b-d** Calnexin intensity and distribution over 10 µm line for each cell. **e-f** Somatic Calnexin area and intensity normalized with MAP2 signals. **g** Representative images of Calreticulin (red) and MAP2 (red) in control and R451C iNs. **h-j** Calreticulin intensity and distribution over 10 µm line for each cell. **k-l** Somatic Calreticulin area and intensity normalized with MAP2. **m** Analysis of ER stress markers (immunoblot) in the excitatory-inhibitory co-cultured iNs. **n** Quantification of ER stress markers protein expression level N=6, 3 cultures 2 repeats. **o** Representative images of Caspase-3 (red), MAP2 (green), and DAPI human neurons carrying wildtype or R451C NLGN3. **p** Quantification of neuronal density (DAPI and MAP) and caspase-3^+^ cells. Statistical significance (^*^p<0.05; ^**^p<0.01) was evaluated with Student’s t-test (bar graphs).

### The *NLGN3* R451C-mutation increases excitatory synaptic connectivity of human neurons transplanted into mouse brain

Cultured neurons lack a physiologically relevant environment. Grafting of human neurons into rodent brains has been used to study human neuronal development and functionality^26, 38^. We transplanted control and *NLGN3* R451C-mutant excitatory human neurons expressing mCherry and mVenus, respectively, together into the hippocampus of neonatal (postnatal day 0-3) RAG2 immunocompromised mice (**Fig. 4a**), and analyzed the neurons six weeks later by imaging and electrophysiology. Human iNs were found to be located mainly at the injection sites but exhibited extensive branching around these sites (**Fig. 4b & c**).

**Fig. 4.**
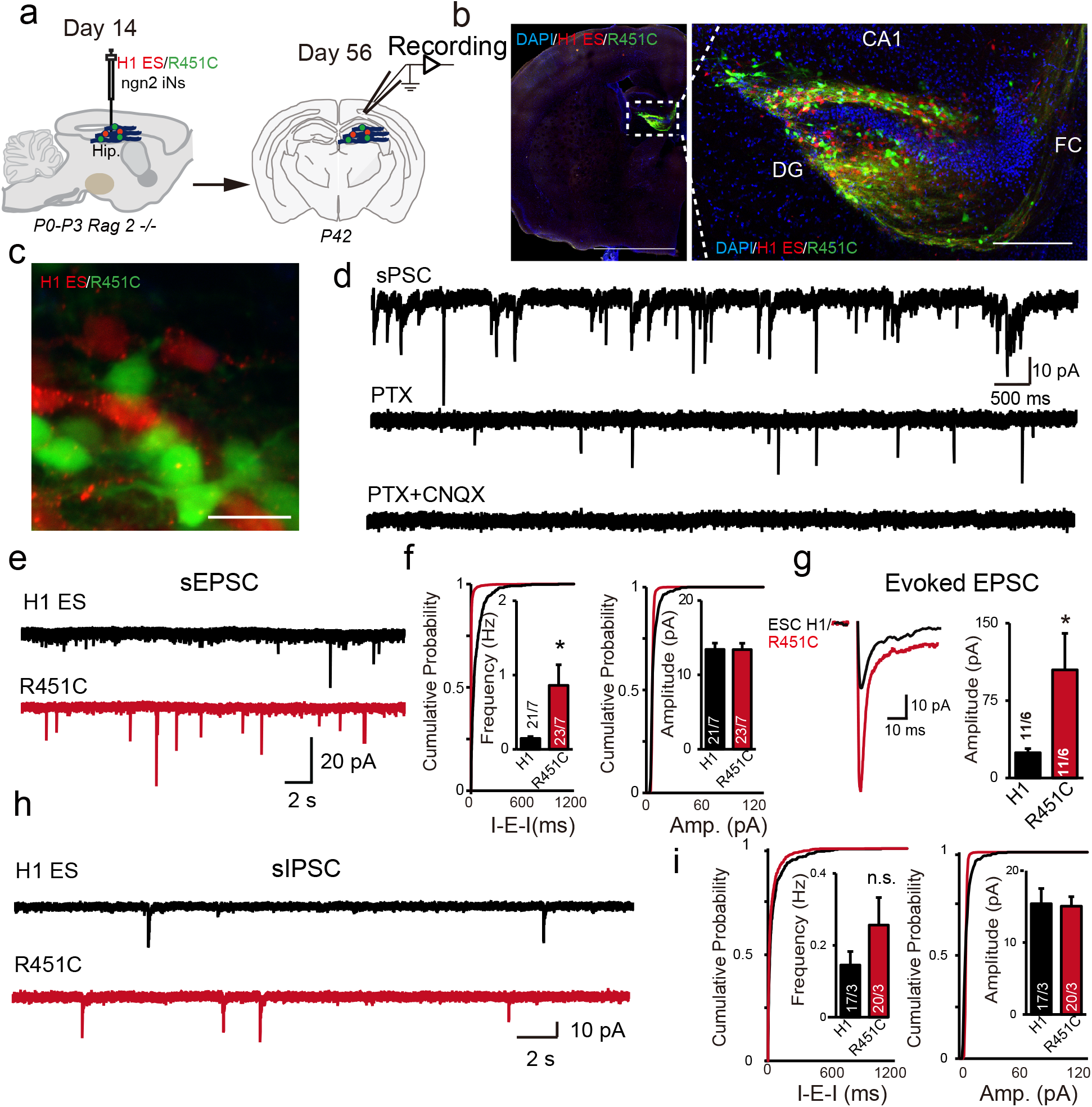
Dual-color genotype transplantation of ESC H1 and NLGN3 R451C human iN cells *in vivo*. **a** Experimental design of control (mCherry) and R451C (GFP) iNs transplant in vivo. **b** Representative images of iN cells transplanted in Rag2-/- hippocampal region, a control line (mCherry), and R451C lines (GFP). **c** Representative images of control and R451C converted iNs in vivo. **d** Representative sPSC trace of transplanted iNs in vivo. Slow responses could be blocked by picrotoxin (PTX). In the presence of PTX and CNQX (both 50 μM), no spontaneous activities were observed. **e** Representative sEPSC trace of control/R451C in vivo. **f** Quantification of sEPSC frequency and amplitude in vivo. **g** Representative trace and quantification of evoked EPSCs. **h** Representative sIPSC trace of control/R451C in vivo. **i** Quantification of sIPSC frequency and amplitude in vivo. The black and red bar represents control ES cell line and R451C converted cell lines. Data are mean + SEM; Number of cultured cells/ culture batches are shown in bars. Statistical significance (^*^, p<0.05) was evaluated with the Kolmogorov-Smirnov test (cumulative probability plots) and Student’s t-test (bar graphs).

We next performed whole-cell patch-clamp recordings from the transplanted neurons in acute hippocampal slices. We recorded from mCherry-positive (control iNs) and mVenus-positive neurons (*NLGN3* R451C-mutant iNs) in the same slices to control for inter-animal variations caused by the transplantation procedure. The transplanted human iNs received both excitatory and inhibitory synaptic inputs (**Fig. 4d**), suggesting that the grafted human neurons were functionally integrated into existing *in vivo* neuronal networks. As usual in such experiments, the integration of the transplanted neurons was only examined postsynaptically, which is appropriate since NLGN3 is a postsynaptic protein. Similar to the findings in in cultured neurons (Fig. 3), *NLGN3* R451C-mutant neurons derived from both ES cell clones exhibited a similar increase in the frequency but not amplitude of sEPSCs (**Fig. 4e-f**). Consistent with the increase in sEPSC frequency, we observed a large increase in evoked EPSC amplitude in *NLGN3* R451C-mutant neurons compared to controls (**Fig. 4g**). We did not find any significant differences between R451C and control neurons in sIPSC frequency or amplitude, but there was a strong trend towards an increase in frequency (**Fig. 4i**). These results confirm that the *NLGN3* R451C-mutation enhances excitatory synaptic connectivity also in an in vivo paradigm, consistent with a gain-of-function phenotype.

### The *NLGN3* R451C-mutation is associated with distinct gene expression profiles in excitatory and inhibitory neurons

To gain insight into the underlying molecular mechanism of R451C *NLGN3* in human neurons and the observed synaptic phenotypes, we conducted single-cell RNA sequencing (sc-RNAseq) experiments in mixed cultures of excitatory or inhibitory neurons derived from control or *NLGN3* R451C-mutant ES cells (**Fig. 5a**). A total of 27,724 cells with high quality and sequencing depth were obtained (average number of genes/cells 3,995; average number of reads 17,574) (**Supplemental Fig. 3**). Cluster identities were estimated using the top cluster-enriched genes. Despite the relatively homogenous human neuronal populations expected from using iN technologies^35-37^, gene expression profiles corresponding to seven cell types were identified: 1) Relatively mature excitatory neurons (*GAP43, DCX, MAP2, NEFM, VMAP2, NSG1, NCAM1, SYT1, STMN2*); 2) Relatively mature NPY inhibitory neurons (*TAC1, NTNG1, NTS, CALB1, NOS1, NPY, GAD2, GAL*), 3) SST inhibitory neuronal cells (*NNAT, SSTR2, CNTN2, SST*); 4) Immature excitatory neurons (*NES, KCNQ1OT1, CACNA1A, DCX, SCN3A, MAP6, NTM, NEFM, NNAT, KCNJ6, NEUROD1, NEUROD4, HOMER3, SLC17A7*); 5) Immature inhibitory neurons (*VIM, NES, DCN*) and 6) Inhibitory neurons (*NNAT, SSTR2, CNTN2, NTRK2, SOX2, CENPF, NES, ID3, TOP2A, PEA15, ID1, HES5, POU5F1, CENPF*); and 7) Neuronal progenitor cells (*SOX2, NES*) (**Fig. 5b, Supplemental Fig. 4 &Table 2**). The distributions of the seven major cell types were reasonably consistent among neurons derived from control ES cells and the two clones of *NLGN3* R451C-mutant ES cells **(Supplemental Fig. 5)**. We observed robust expression of genes that characterize human cortical layers I-V (*RELN*, Layer I; *CUX1, POU3F2*, layer II-III; *NECAB1*, layer IV and *BCL11B*, layer V) in induced human iNs, suggesting a phenotype reminiscent of cortical identity (**Fig. 5 c**)^36, 37^.

**Fig. 5.**
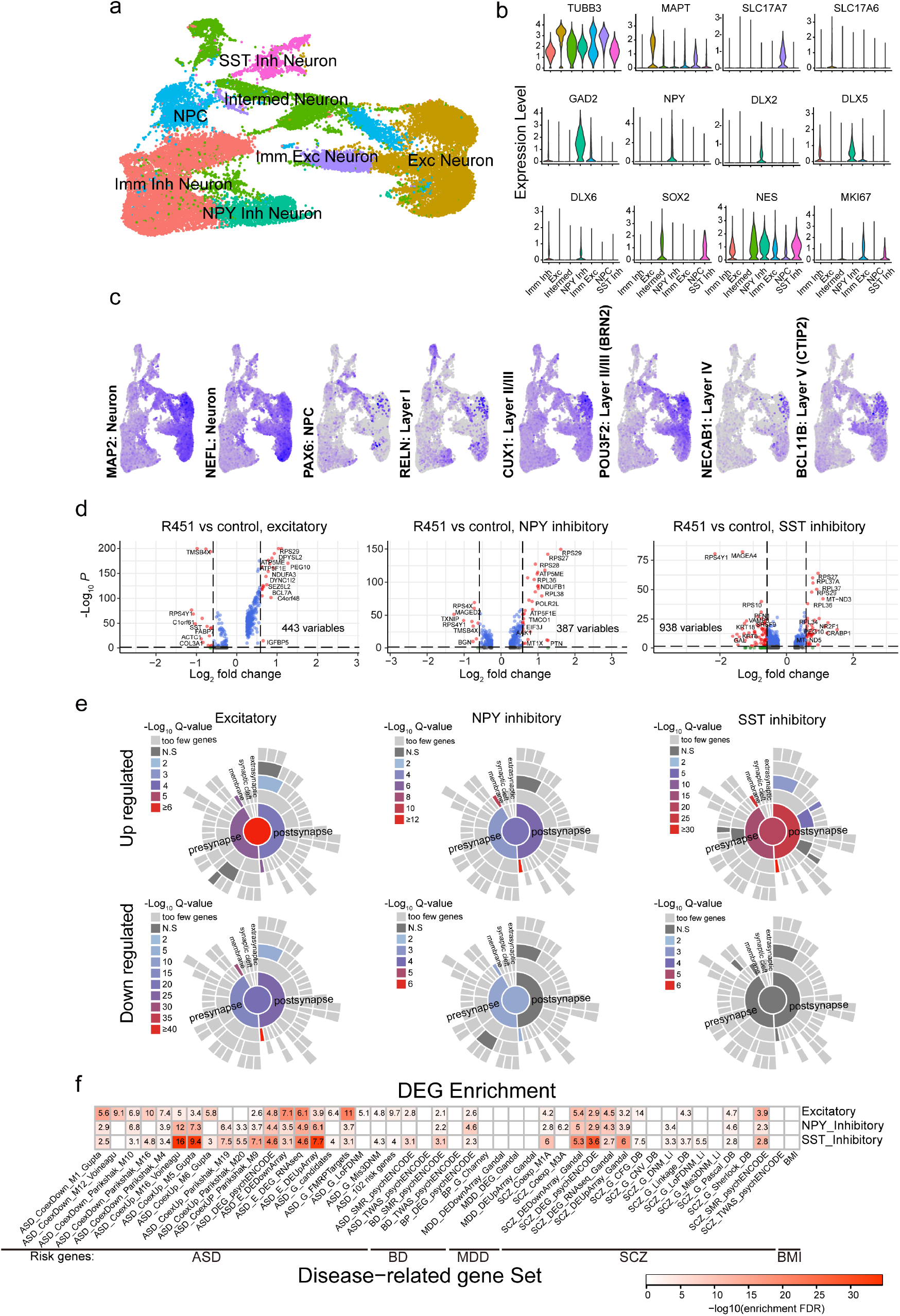
Single-cell RNA seq of E/I mixed culture R451C human iNs. **a** UMAP of all human neuronal cells (n=27,724) reveals seven distinct neuronal subtypes. **b** Expression levels of human neuron and progenitor marker for each neuronal subtype. **c** Human induced neurons express mature neuron and cortical layer marker features. **d** DEGs in R451C versus control in excitatory, NPY and SST inhibitory neuron subtypes with all replicates pooled. Differential expression is defined by FDR<0.05 and log2FC>0.25. **e** Sunburst plots of enriched synGO term in excitatory, NPY, and SST inhibitory neuron subtypes. **f** Overrepresentation of ASD, SCZ, BD, MDD, and BMI-related genes in excitatory, NPY, and SST inhibitory neuron subtypes DEGs. The color of the box shows the odds ratio for enrichment (shades of red for significant enrichment, adjust p<0.05).

To investigate the potential impact of the *NLGN3* R451C-mutation on gene expression in neuronal cell clusters associated with synaptic phenotypes, we selected the relatively mature excitatory cluster and the inhibitory clusters (both NPY and SST) for analyses. We examined the differentially expressed genes (DEGs; false discovery rate (FDR) < 0.05 and log_2_ fold change [log2FC] > 0.25) within individual clusters, comparing *NLGN3* R451C-mutant and control iNs (**Supplemental Table 3**). We identified 443 genes as DEGs in the excitatory cluster, 387 DEGs in the NPY inhibitory cluster, and 938 DEGs in the SST cluster. Interestingly, these DEGs were mostly unique to the selected neuronal subtypes (26.5%-1.9% overlaps), suggesting that the *NLGN3* R451C-mutation may have distinct effects in different types of neurons (**Fig. 5d**). For example, the ASD-related gene *FXR1* is upregulated in *NLGN3* R451C-mutant excitatory neurons (adjusted *P=*2.95 ×10^−58^) and NPY inhibitory neurons (adjusted *P=*2.75× 10^−18^), but not in SST inhibitory neurons.

We then applied SynGo annotations to all three neuronal type clusters using isolated upregulated and downregulated gene sets^49^. We found three groups of upregulated genes that are enriched in genes related to pre- and postsynaptic function categories. However, the excitatory neuron cluster exhibited a stronger enrichment in downregulated synaptic-related genes, especially postsynaptic functions (**Fig. 5e**). These annotations are consistent with the idea that the *NLGN3* R451C-mutation plays heterogeneous roles in different neuronal cell types. The excitatory neurons show more profound DEGs that are associated with synaptic transmission, for example, the synaptic vesicle exocytosis genes *VAMP2, SV2A, ATP6V0C, ATP6V0A1* are all upregulated in R451C *NGLN3* excitatory neurons but not in inhibitory neurons (**Supplemental Table S4**). These results suggest that *NLGN3* R451C-mutation may have a more profound impact on excitatory synaptic transmission, consistent with our observations using *in vitro* and *in vivo* human neuronal models. However, it should be noted that these comparisons do not analyze truly isogenic cells, but neurons derived from a parental ES cell line with neurons derived from mutant clonal ES cell lines that were generated from the parental line. Since clonal derivative of human stem cell lines can exhibit large changes in gene expression simply by the effect of their clonal propagation^28^, it is unclear whether these gene expression changes are due to the *NLGN3* R451C-mutation or to the clonal derivation of the ES cells.

To further investigate the interaction between *NLGN3* and DEGs, a protein-protein interactions (PPIs) network associated with NLGN3 was constructed with *NLGN3* and DEGs identified in three neuronal clusters (excitatory neurons, NPY, and SST inhibitory neurons). The first- and second-level neighbors to *NLGN3* were shown in **Supplemental Fig. 7**. Notably, we found a correlation of the *NLGN3* R451C-mutation with CASK expression. *CASK* is a cytoplasmic scaffolding protein that interacts with neurexins that bind to NLGN3 and that are associated with ASDs^62-65^. *CASK* protein levels but not RNA expression is increased by a heterozygous conditional *NRXN1* deletion of human neurons, which exhibit a decrease in excitatory synaptic strength^29^. *CASK* deletion in mice, conversely, results in an increased mESPC frequency in cortical neurons^66^.

To determine whether *NLGN3* affects the connection between neurons, we performed cell-cell interaction analysis in the R451C dataset and control dataset ^50^. We calculated the cell-cell interaction based on the expression of known ligands and receptors in cell clusters. Interestingly, we found an overall increase in cell-cell interaction number and strength in R451C neurons compared to the control dataset (**Supplemental Fig. 6 a&b**). To identify the molecular pathway mediating the increase in interaction, we aggregated the interactions of ligand-receptor pairs by pathways. In the excitatory neuronal cluster, we found a significant increase in interaction in *NLGN3* binding partner *NRXN1*, as well as cell adhesion related *NEGR* that may enhance synaptic function (**Supplemental Fig.6c**)^7^. Thus, the concerted upregulation of these binding pairs in R451C excitatory neurons again strongly suggests a role for R451C function in enhancing synaptic transmission.

Finally, to further determine whether the DEGs associated with the *NLGN3* R451C-mutation are related to neuropsychiatric disorders such as ASDs, schizophrenia (SCZ), bipolar disorder (BP), and major depression disorder (MDD), we performed an enrichment test in the risk genes of ASD, SCZ, BP, and MDD using body mass index (BMI) as a control. The risk genes being tested were defined as genes showing genetic association, differential expression, or co-expression in psychiatric patient postmortem brains^67-85^ (**Fig. 5f, Supplemental Table 5-7**). We observed significant enrichment in DEGs of all three cell types (excitatory, NPY, and SST inhibitory neuron clusters) with ASD, SCZ, and BP-related gene sets. However, MDD and BMI did not show significant enrichment in the analysis. In excitatory neurons we found that 13 genes are shared in signals of genetics, DEGs, and COE level (*VAMP2, DDX1, KIF1A, KLC1, PPP6R2, ATP6V0A1, SV2A, AAK1, KIF1B, SLC8A1, SEZ6L2, ANK3, DDX24*). In NPY neurons, 5 genes are shared in signals of genetics, DEGs, and COE level (*DDX1, PPP6R2, LTBP1, AAK1, SEZ6L2*). In SST neurons 10 genes are shared (*CARHSP1, TTC3, NREP, CPE, FABP5, CKB, PPP2R1A, AP2S1, S100A10, CADM1*) (**Supplemental Fig. 8**). These results suggest that convergent gene networks that are modified by the *NLGN3* R451C-mutation play pivotal roles in the pathophysiology of mental disorders including SCZ, BP, and ASD.

## DISCUSSION

Here we show that the R451C-mutation of *NLGN3*, the first mutation identified in an ASD patient^8^, increases excitatory, but not inhibitory, synaptic connectivity in human neurons. This phenotype was observed in human neurons both when analyzed in culture and when examined after transplantation into mouse brain. Analyses of other neuronal parameters, such as neuronal dendrites or intrinsic electrical properties, uncovered no changes. We found that despite the fact that the R451C-mutation decreases the levels of *NLGN3* protein, the increase in excitatory synaptic connectivity induced by this mutation is not associated with an ER stress response, but represents a gain-of-function phenotype. Moreover, we describe that *NLGN3* R451C-mutant neurons exhibit major gene expression changes when compared to neurons derived from their parental ES cell line. Our findings thus show that in human neurons, the *NLGN3* R451C-mutation causes a selective increase in excitatory synaptic function, suggesting a potential pathophysiological mechanism by which the R451C-mutation induces ASDs.

*NLGN3* is an adhesion molecule that is essential in mice for the normal function but not the formation of both excitatory and inhibitory synapses^59, 86-88^. Mutations in *NLGN* genes have been shown to induce ASDs and intellectual disability with an apparently 100% penetrance in multiple families^89-92^. In contrast, mutations in neurexins, especially *NRXN1*, predispose to a larger spectrum of neurodevelopmental disorders with a lower penetrance^28, 29, 93-95^.

Although the *NLGN3* R451C-mutation was intensely studied in knockin mice, little was known about its effect in human neurons. In the mice, the *Nlgn3* R451C-knockin mutation recapitulated, at least in part, the cognitive and motor behavior alterations of ASD subjects^12, 15, 16, 96, 97^. A series of detailed analyses in mice revealed that the R451C-mutation induces region-, cell type-, and synapse-specific changes at both excitatory and inhibitory synapses that are consistent with gain-of-function and loss-of-function mechanisms at different synapses^12-14, 19^. For example, the *Nlgn3* R451C-mutation enhanced inhibitory synaptic transmission in the cortex^12^, but suppressed parvalbumin-positive inhibitory synapses in the hippocampus^19^. Furthermore, excitatory synaptic transmission was enhanced in hippocampal neurons and in cerebellar Purkinje cells^13, 14^. These data supported the hypothesis that *NLGN3* acts in specifying synapse properties in a manner that depends on the synaptic contact, suggesting that presynaptic *NLGN3*-binding partner that are differentially expressed in different synapses may be crucial determinants of its role^7^.

Extensive studies on the effect of the *Nlgn3* R451C-mutation on synapses in mice have strongly supported a causative role for the R451C-mutation in ASD pathophysiology, but it was unclear from these studies whether the results in mice translate to human neurons. Studies of human neurons carrying ASD-linked mutations in *NRXN*1^28, 29^ or *NLGN*4X^30, 31^ uncovered robust human-specific synaptic phenotypes, suggesting that the *NRXN*-*NLGN* adhesion complex may perform different roles in mice and humans. For example, heterozygous *NRXN1* deletions produced profound synaptic deficits in human but not in mouse neurons that were generated by identical procedures^28, 29^. The goal of the current study therefore was to examine whether the *NLGN3* R451C-mutation alters the function of synapses in human neurons similar to mouse neurons.

We found that in human neurons, the *NLGN3* R451C-mutation augmented the strength of neurotransmission and increased the overall number of excitatory synapses (a gain-of-function phenotype), but had no major effect on inhibitory synapses (**Fig. 1 & 3**). Importantly, the hyper-glutamatergic transmission phenotype was maintained in *NLGN3* R451C-mutant neurons after they had been grafted into mouse brain (**Fig. 4**). Given the cell type- and region-specific effects of the mouse *Nlgn3* R451C-mutation as discussed above, it is important to note that other types of human neurons might need to be considered to fully assess the impact of the *NLGN3* R451C-mutation on the human brain. Nevertheless, the current experiments establish that the observation of a gain-of-function effect of the *Nlgn3* R451C-mutation in mouse neurons is transferable to human neurons, and that the *NLGN3* R451C-mutation produces a profound and selective change in synaptic function in human neurons that could reasonably account for ASD pathogenesis.

Using single-cell gene expression profiling, we found that the *NLGN3* R451C-mutation is associated with differential gene expression changes in excitatory and inhibitory human neurons. Excitatory *NLGN3* R451C-mutant neurons showed a higher synaptic function enrichment than inhibitory *NLGN3* R451C-mutant neurons. Furthermore, we found an upregulation of the presynaptic *NLGN3* binding partner *NRXN1*, which may enhance synaptic function specifically in excitatory neurons. We conducted a comprehensive comparison between R451C DEGs and psychiatric disorders’ risk genes, taking advantage of numerous types of data, including genetic findings, differential expression, and co-expression studies (**Supplemental Table 2**)^67-85^ to capture different aspects of genetic, environmental, or gene– environmental interaction effects. Analysis of the gene network of the DEGs of R451C-mutant neurons suggested that they associated strongly with ASD, SCZ, and BP but not with MDD and obesity. These data imply that different mental disorders may have convergent gene networks that affect neuronal functions including synaptic transmission in the brain^98^. We hypothesize that autism may be associated with dysregulated convergent gene networks, and ASD-related gene mutations in a single gene, such as *NLGN3*, may affect this convergent gene network function.

As any study, our experiments have inherent limitations. It has been shown that derivation of genetically altered cell clones from human stem cells causes a profound change in gene expression because stem cells are undergoing continuous genetic changes in culture that are often subtle but accumulate during clonal expansion^28^. As a result, the RNAseq profiles of neurons derived from a stem cell line and from wild-type clonal derivatives of that stem cell line differ dramatically^28^. Thus, the assessment of DEGs between neurons generated from parental stem cell lines and clonal mutant derivatives needs to be treated with caution, as these neurons are not truly isogenic and isogenicity can only be achieved with conditional manipulations. However, the fact that the DEGs we observed in our study between R451C-mutant and control neurons are consistent with a change in synaptic pathways is encouraging a biological interpretation, although such an interpretation will remain subject to reservations until isogenic neurons are examined. It should be noted that this is a general problem in studies of gene expression changes of mutant neurons, and that a precise set of comparisons is often difficult to achieve.

Another limitation of our study is that we only considered a limited number of excitatory and inhibitory neuronal types, and also did not examine any glial cell types. *NLGN3* is also expressed in glial cells (**Supplemental Fig. S1e**)^55, 56^. The effect of the *NLGN3* R451C-mutation on glial cell types and different brain regional human neurons require further investigations.

Furthermore, we have not explored the trafficking and expression of NLGN3 R451C-mutant protein in the human neurons, although extensive studies on this question were performed previously using heterologous expression approaches^10^. Nevertheless, we have explored the hypothesis that ER stress and UPR may be involved in the pathogenesis associated with R451C^14^ but found inconsistent results (**Fig. 3**).

In summary, the present study dissected the functional impact of the autism-related *NLGN3* R451C-mutation on synaptic transmission in human neurons. The dramatic enhancement of excitatory synaptic connectivity by the *NLGN3* R451C-mutation suggests a pathophysiological pathway in the development of ASDs whereby increases in cortical glutamatergic synaptic transmission may alter the overall excitatory/inhibitory balance, consistent with earlier hypotheses that changes in this balance are a major driver of neuropsychiatric disorders.

## Acknowledgments

We thank the members of the Pang Lab for their insightful comments on the manuscript. We also thank Xueying Wang from Central South University for collecting the psychiatric risk summary gene list.

## Author contributions

L.W. conducted the experiments including cell culture, electrophysiology, morphological and genomics analysis. V.R.M. conducted the CRISPR/Cas9 gene targeting, related analyses and provided conceptual input on experimental design. R.D., X.S., R. X., M.B., I.S., Y.C., J.T., conducted part of the analysis. R.H. conducted the single cell analysis. K.Y.K, P.J., D.C., C.P., T.C.S, C.L., C.C. planned the experiments and analyzed the data. V.R.M., L.W., D.C., Z.P.P. and T.C.S. conceived the project. T.C.S. and Z.P.P. wrote the paper with input from all authors.

